# SID-2 localises to extracellular vesicles in parasitic nematodes and does not function in environmental RNAi

**DOI:** 10.1101/2024.07.03.599821

**Authors:** Frances Blow, Kate Jeffrey, Franklin Wang-Ngai Chow, Inna A. Nikonorova, Maureen M. Barr, Atlanta G. Cook, Bram Prevo, Dhanya K. Cheerambathur, Amy H. Buck

## Abstract

In the free-living nematode *Caenorhabditis elegans* the transmembrane protein SID-2 imports double-stranded RNA into intestinal cells to trigger systemic RNA interference (RNAi), allowing organisms to sense and respond to environmental cues such as the presence of pathogens. This process, known as environmental RNAi, has not been observed in the most closely related parasites that are also within clade V. Previous sequence-based searches failed to identify *sid-2* orthologs in available clade V parasite genomes. In this study we identified *sid-2* orthologues in these parasites using genome synteny and protein structure-based comparison, following identification of a SID-2 orthologue in extracellular vesicles from the murine intestinal parasitic nematode *Heligmosomoides bakeri*. Expression of GFP-tagged *H. bakeri* SID-2 in *C. elegans* showed comparable localisation to the intestinal apical membrane as seen for GFP-tagged *C. elegans* SID-2 and further showed mobility in intestinal cells in vesicle-like structures. We tested the capacity of *H. bakeri* SID-2 to functionally complement environmental RNAi in a *C. elegans* SID-2 null mutant and show that *H. bakeri* SID-2 does not rescue the phenotype in this context. Therefore, our work identifies SID-2 as a highly abundant nematode EV protein whose ancestral function may be unrelated to environmental RNA and rather highlights an association with extracellular vesicle-mediated communication in free-living and parasitic nematodes.

## BACKGROUND

Uptake of exogenous double-stranded RNA (dsRNA) and subsequent entry into RNA interference (RNAi) pathways enables organisms to sense and respond to environmental cues and is a potent mechanism of antiviral defence in some species [1]. Environmental RNAi has been characterised in the model nematode *Caenorhabditis elegans* using genetic screens [2,3] and the gene systemic RNA interference defective 2 (SID-2) was discovered as being essential for dsRNA import [4]. SID-2 is a single-pass transmembrane protein that is predominantly localised to the apical membrane of intestinal epithelial cells and is thought to play a role in the internalisation of dsRNA through pH-dependent receptor-mediated endocytosis [5]. However, the precise mechanism by which SID-2 mediates dsRNA import and entry into RNAi pathways remains unknown. Although *sid-2* orthologues have been identified in many *Caenorhabditis* species, most are not capable of environmental RNAi, including one of the closest known phylogenetic relatives of *C. elegans*, *C. briggsae* [4,6].

Nematodes are an incredibly diverse and ubiquitous phylum that includes free-living species and species that parasitise plants, animals and humans [7]. *Heligmosomoides bakeri* is a murine gastrointestinal nematode that naturally infects house mice (*Mus musculus*) and belongs to the same phylogenetic clade as *C. elegans* (clade V) which also includes parasites infecting livestock and humans [8]. *H. bakeri* has been used as a laboratory model to study parasite manipulation of the host immune system due to its ability to cause chronic infections in mice via the secretion of immunomodulatory molecules [9]. Like most animal parasitic nematodes, *H. bakeri* is refractory to environmental RNAi [10]. This was previously hypothesised to be at least partially due to a lack of *sid-2* orthologues in this group of species [11,12]. We report here for the first time that *C. elegans sid-2* orthologues were identified in clade V parasitic nematodes using genome synteny and protein structure comparison tools in conjunction with chromosome-scaffolded genome assemblies. A phylogenetic analysis of *sid-2* orthologues from free-living and parasitic clade V nematodes revealed that the extracellular domain of SID-2 is highly divergent between species, in both free-living and parasitic nematodes.

In parallel, a recent study suggests SID-2 may play roles in other contexts, since it is released as an abundant cargo of ciliary extracellular vesicles (EVs) by *C. elegans* [13]. Previous studies have demonstrated a role of EVs released by *C.elegans* in worm-to-worm communication via manipulation of mating behaviours [13,14]. One way in which *H. bakeri* and other nematode parasites manipulate the host immune system is through the secretion of EVs bearing immunomodulatory cargo including small RNAs, which are taken up by host cells [15–18]. Based on a previous proteomic analysis, we found that, like *C. elegans* SID-2, *H. bakeri* SID-2 is abundant in EVs. This suggests that the conservation of SID-2 could relate to functions in EVs rather than environmental RNAi. We investigated the function of the SID-2 orthologue from the parasitic nematode *H. bakeri* by heterologous expression in *C. elegans*. We found that *H. bakeri* SID-2 does not compensate for the *C. elegans* SID-2 environmental RNAi phenotype under endogenous *C. elegans* regulatory elements, indicating that the protein function at the intestinal luminal membrane is divergent between *C. elegans* and *H. bakeri* SID-2 orthologues. This work expands the contexts in which SID-2 exists and introduces the question of whether an ancestral function of nematode SID-2 proteins could relate to cell-cell, nematode-nematode or nematode-host communication via EVs.

## RESULTS

### a. Identification of an *H. bakeri* protein orthologous to *C. elegans* SID-2 that is enriched in extracellular vesicles

A previous proteomic analysis identified proteins enriched in EVs compared to EV-depleted *Heligmosomoides* excretory-secretory products (HES) from *in vitro* cultured *H. bakeri* adult worms [15]. Several of these EV-enriched proteins have no annotated function (**Table S1**). One of the abundant EV proteins, HPOL_0001199201 (WormBase ParaSite accession), which exhibits a 17.5-fold enrichment in EV fractions compared to non-EV fractions in the proteomic analysis (**Fig. 1A**; **Table S1**), was analysed using BLASTp [19]. The search was conducted against *C. elegans* annotations in WormBase and identified SID-2 (protein accession ZK520.2) as a potential orthologue.

**Figure 1.**
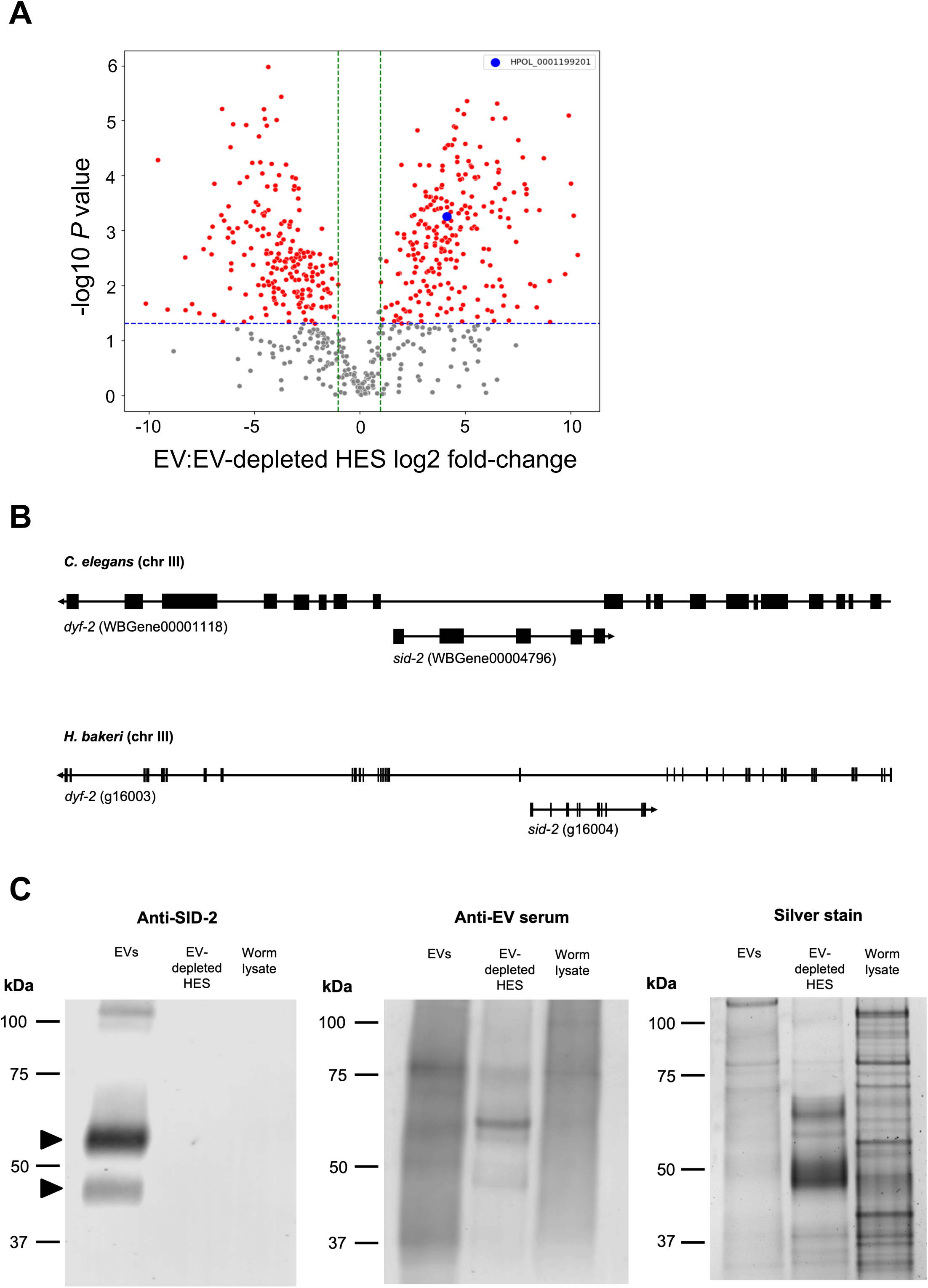
The parasitic nematode *H. bakeri* encodes a direct orthologue of *C. elegans sid-2* that is enriched in extracellular vesicles released by the parasite. **A** Scatter plot of proteins enriched in EVs compared to EV-depleted HES identified by LC-MS/MS from [15]. The *sid-2* orthologue, HPOL_0001199201, is highlighted in blue. Red points indicate *p* < 0.05 and log2 fold change (FC) > 1, while grey points indicate non-significant proteins. The blue horizontal dashed line represents the *p*-value threshold of 0.05, and the green vertical dashed lines represent log2 FC thresholds of 1 and −1. **B** Genome synteny of *sid-2* orthologues on chromosome III in *C. elegans* and *H. bakeri*. In both genomes, *sid-2* is encoded on the opposite strand in an intron of a larger gene *dyf-2*. Black boxes indicate exons, and genes are displayed from start codon to stop codon. Arrows indicate gene orientation in the genome. **C** Equal amount of total protein (1 μg) isolated from *H. bakeri* adult worm lysate, EVs, and EV-depleted HES was assayed by anti-SID-2 (left panel) and anti-EV serum (middle panel) western blot and silver stain (right panel). SID-2 bands ∼45 kDa and ∼55 kDa enriched in EVs are marked with black triangles on the anti-SID-2 western blot (left panel).

Because genome synteny is highly conserved in clade V nematodes [22], we compared the genomic locations of *C. elegans sid-2* and *H. bakeri* HPOL_0001199201. In *C. elegans* genome assembly WBcel235 on WormBase (version WS290) *sid-2* (WBGene00004796) is nested within the larger gene *dyf-2* (WBGene00001118) on chromosome III (**Table 1**, **Fig. 1B)**. Similarly, HPOL_0001199201 (annotated as g16004 in an updated chromosome-scale genome assembly for *H. bakeri* nxHelBake1.1 [20]) is nested within a larger gene *dyf-2*-like (g16003) on chromosome III (**Table 1**, **Fig. 1B**). Amino acid identity between *C. elegans* SID-2 and HPOL_0001199201 is only 24.5 % (**Fig. S1A**). However, similarities in the AlphaFold models generated using AlphaFold 3 [23] for HPOL_0001199201 (**Fig. S1B**) and *C. elegans* SID-2 (**Fig. S1C**) indicate structural similarity such as the conserved barrel in the extracellular domain of both proteins, the single-pass transmembrane alpha helix, and the disordered cytoplasmic domain, although low confidence for some regions of the models. Similarly, a FoldSeek [24] search of the *C. elegans* SID-2 AlphaFold model against the AlphaFold database restricted to *H. bakeri* proteins returns HPOL_0001199201 as the top hit, with a probability of 1.0. Given the shared genome synteny and protein structural similarity of HPOL_0001199201 and *C. elegans sid-2*, we hereafter refer to HPOL_0001199201 as *H. bakeri sid-2*.

**Table 1.** Details of genomic loci encoding *H. bakeri* and *C. elegans sid-2* orthologues.

| Organism | Genome assembly name | Gene accession | Chromosome | Gene start coordinates | Gene end coordinates | Citation |
| --- | --- | --- | --- | --- | --- | --- |
| *H. bakeri* | nxHelBake1.1 | g16004 | III | 13899061 | 13907575 | [20] |
| *C. elegans* | WBcel235<br>version WS290 | WBGene00004796 | III | 13679445 | 13682450 | [21] |

Consistent with the proteomic analysis, western blot analysis using an antibody raised against *H. bakeri* SID-2 confirms enrichment of SID-2 in EVs when loading equal amounts (1 μg) of total protein from *H. bakeri* adult worm lysate, EVs (approx. 2.4 x 10^9^ particles loaded), and EV-depleted HES (**Fig. 1C**; left panel). *H. bakeri* SID-2 has a predicted molecular weight of 39 kDa but migrates at ∼45 kDa and ∼55 kDa (marked with black triangles) in the EV sample, with neither band observed in EV-depleted HES or worm lysate. The disparity in predicted and observed molecular weight may relate to modifications such as glycosylation (*H.bakeri* SID-2 is predicted to have three N-glycosylation sites by NetNGlyc version 1.0) and/or altered migration as a transmembrane protein [25,26]. The same blot was probed with serum from rats vaccinated with EVs (middle panel), which shows detection of proteins in all three samples (**Fig. 1C**) and highlights in comparison the specificity of SID-2 and SID-2 antibodies for EVs. Silver stain (**Fig. 1C** right panel) further confirms that worm lysate, EV, and EV-depleted HES had sufficient protein loaded.

### b. Clade V nematode parasites encode direct orthologues of *C. elegans sid-2*

Previous homology-based searches failed to detect orthologues of *C. elegans sid-2* in clade V parasitic nematodes [12]. Using the conserved synteny of *sid-2* nested within *dyf-2* (**Fig. 1B**) and recently improved genomes for clade V parasitic nematodes, we identified orthologues of *H. bakeri sid-2* in clade V parasitic nematodes using the ‘Orthologues’ feature in WormBase ParaSite and the gene finding tool miniprot version 0.13 [27] (**Table S2**). Orthologues of *C. elegans sid-2* were identified in *Caenorhabditis* nematodes using the ‘Orthologues’ function in WormBase (**Table S2**). ConSurf analysis [28,29] of all 28 amino acid sequences from both groups was performed on the multiple sequence alignment (**File S1**) to identify regions of the protein that are conserved in clade V nematodes. This indicates that the cytoplasmic domain of *sid*-2 orthologues is more highly conserved than the extracellular domain, and that this is not unique to parasites, but is seen across all clade V nematode *sid-2* orthologues studied (**Fig. 2, Fig. S2B**).

**Figure 2.**
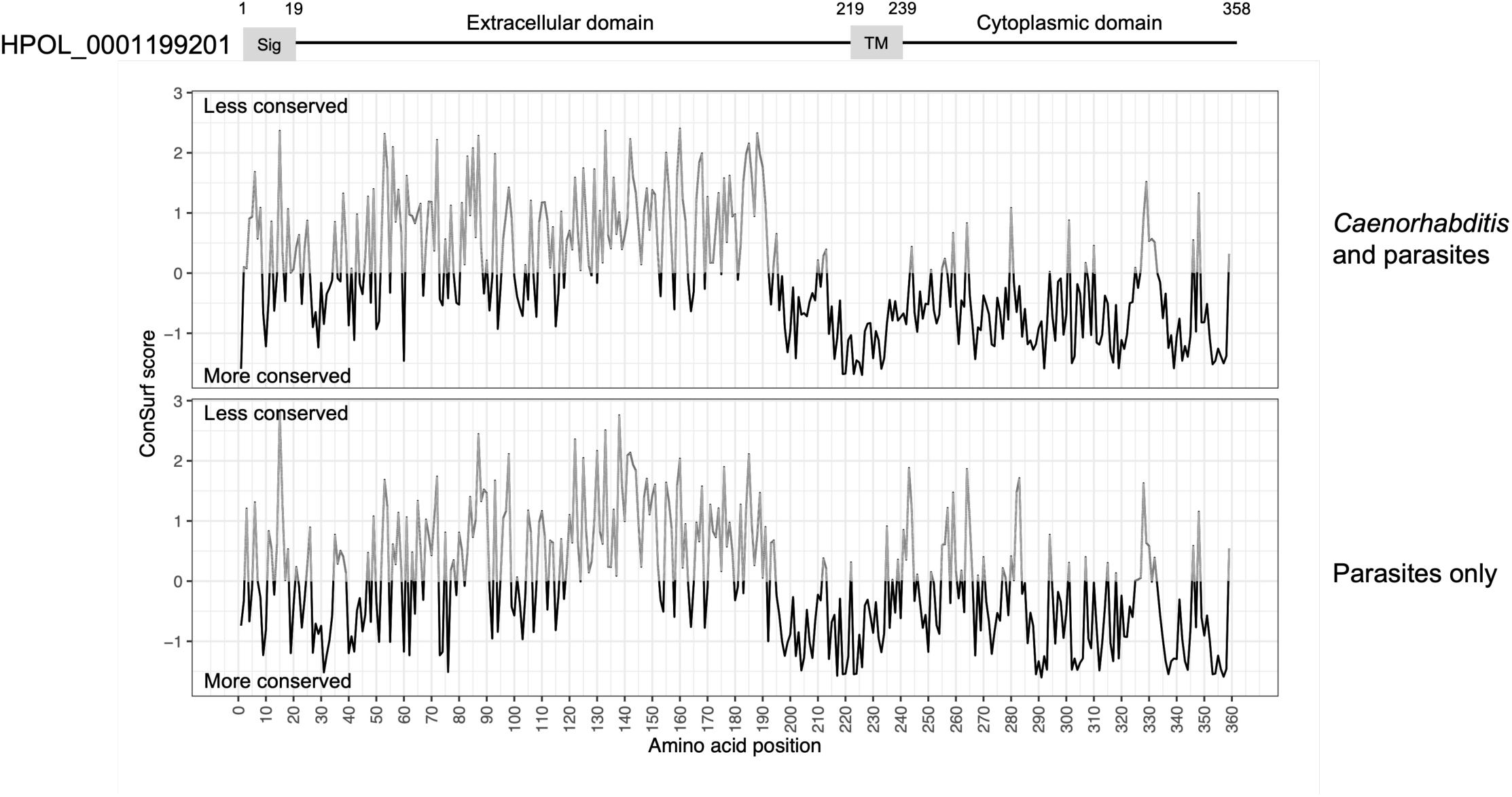
ConSurf analysis of amino acid conservation of HPOL_0001199201 with *sid-2* orthologues from 28 species of clade V nematode including both *Caenorhabditis* and parasitic species (top) and just parasitic species (bottom). Negative values (black) indicate residue conservation and positive values (grey) indicate residue variability at that position.

### c. *H. bakeri* SID-2 localises to the intestinal apical membrane and mobile vesicle-like structures when expressed in *C.elegans*

To test the ability of *H. bakeri* SID-2 to compensate the function of *C. elegans* SID-2 in environmental RNAi, we generated a transgenic *C. elegans* SID-2 null strain expressing a single copy of N-terminally GFP-tagged *H. bakeri* SID-2 under control of the *C. elegans sid-2* promoter (HbSID-2, strain DKC1285). As a positive control, we generated a SID-2 null strain expressing a single copy of N-terminally GFP-tagged *C. elegans* SID-2 under control of the *C. elegans sid-2* promoter using the same method (CeSID-2, strain DKC1365). Imaging of the intestine of the transgenic strains shows similar subcellular localisation between the *H. bakeri* and *C. elegans* SID-2 proteins, localising to the apical membrane of all intestinal cells (**Fig. 3A**) and when zoomed-in GFP-fluorescence appears as vesicle-like structures (**Fig. 3B)**. Imaging of the wild-type N2 strain showed some autofluorescence in the intestine, as expected, but not at the apical luminal membrane (**Fig. 3A**). The expression in both strains is comparable to the apical membrane localisation seen in *C. elegans* with fluorescent tags fused to SID-2 in the endogenous locus via CRISPR-Cas9 modification [13], indicating that transgenic proteins are expressed similarly to endogenous *C. elegans* SID-2.

**Figure 3.**
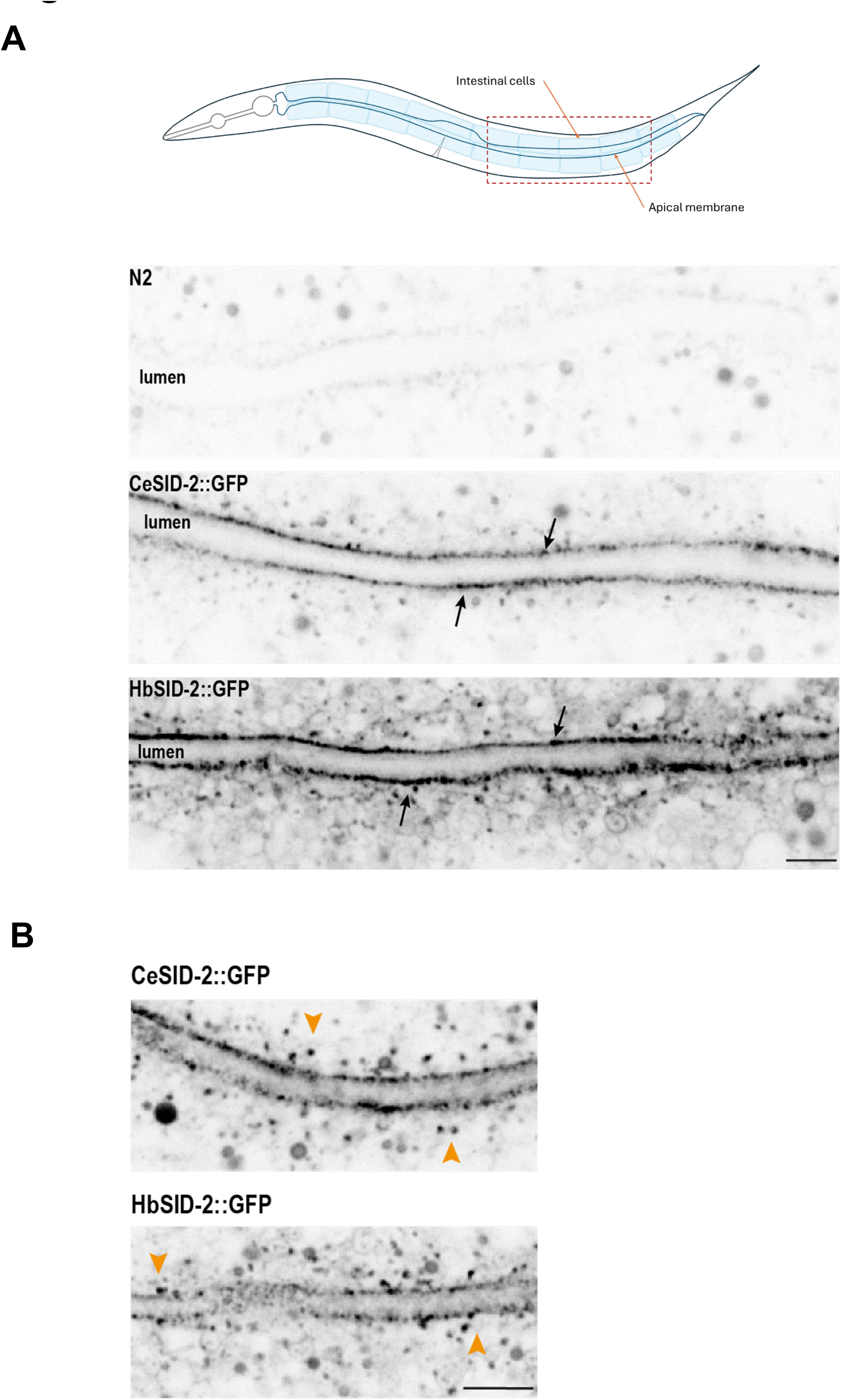
Expression of SID-2::GFP transgenes. **A** (Top) Schematic of the intestine in *C. elegans*. (Bottom) Images of *C. elegans* intestine in N2 (control) animals and animals expressing CeSID-2::GFP and HbSID-2::GFP transgenes. The black arrows point to the apical surface of the intestine. Scale bar, 5 μm. **B** Apical surface view of the *C. elegans* intestine. Orange arrows point to vesicle-like structures budding off from the apical surface in animals expressing CeSID-2::GFP and HbSID-2::GFP transgenes. Scale bar, 2.5 μm.

Corroborating previous evidence that SID-2 is localised to EVs in *C. elegans*, we observed GFP fluorescence and movement consistent with vesicle trafficking in the intestine. This was evident for transgenic strains expressing either *C. elegans* (**Movie S1**) or *H. bakeri* (**Movie S2**) SID-2, suggesting that properties of SID-2 dictating its localisation and mobility are conserved in *C. elegans* and *H. bakeri*.

### d. *H. bakeri* SID-2 does not compensate for *C. elegans* SID-2 function in environmental RNAi

To test the capacity of *H. bakeri* SID-2 to compensate for *C. elegans* SID-2 function in environmental RNAi, we performed an embryonic lethal RNA interference assay. We exposed wild-type N2, SID-2 null, HbSID-2 and CeSID-2 strains to *Escherichia coli* OP-50 expressing dsRNA against *C. elegans* holocentric protein-4 (*hcp-4*), which performs an essential function in chromosome segregation during mitosis [30] and causes embryonic lethality when knocked-down by RNAi [31] (**Fig. 4A**). Offspring viability was significantly different between strains (Kruskal-Wallis = 46.95, p < 0.0001, d.f. = 3), with Dunn’s post-hoc tests indicating significant differences between strains with *C. elegans* SID-2 (wild-type N2 and CeSID-2) and without *C. elegans* SID-2 (SID-2 null and HbSID-2) (**Fig. 4B**). As expected, the proportion of viable offspring was 0% in wild-type N2 and CeSID-2 transgenic worms, consistent with functional uptake of lethal dsRNA from the intestinal lumen by SID-2, and subsequent entry into RNAi pathways in the worm. In the SID-2 null mutant the median proportion of viable offspring was 91.6%, indicating significantly reduced uptake of exogenous dsRNA similar to what was seen with HbSID-2 transgenic worms (98.1%). These results demonstrate that *H. bakeri* SID-2 does not internalise dsRNA from the intestinal lumen, or that internalised dsRNA does not enter functional RNAi pathways (**Fig. 4B**).

**Figure 4.**
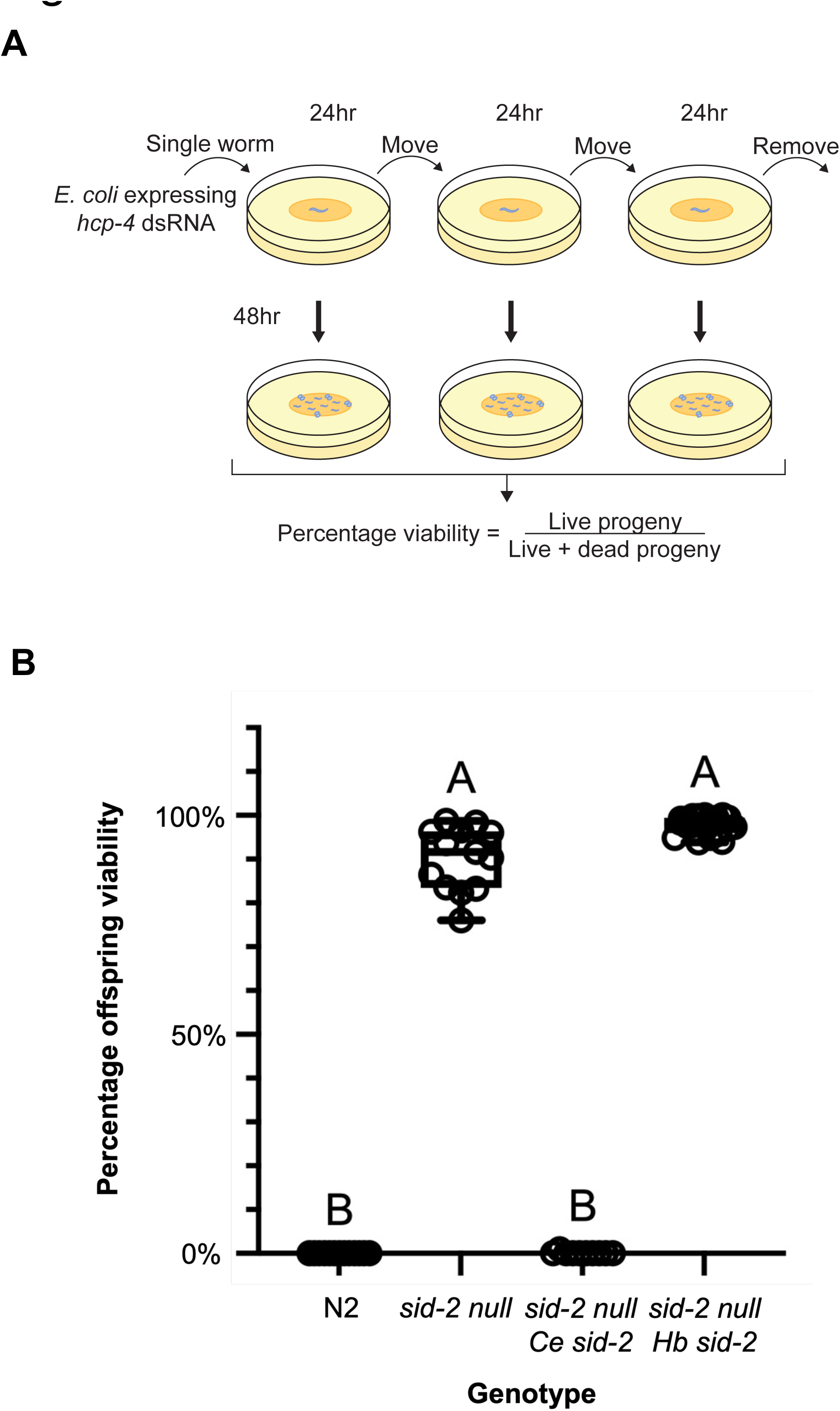
Assay for functional complementation of *C. elegans* SID-2 function in environmental RNAi. **A** Schematic of the environmental RNAi assay. **B** Percentage viability of offspring from SID-2 transgenic worms fed on bacteria expressing embryonic lethal dsRNA against *hcp-4*.

## DISCUSSION

Despite a lack of environmental RNAi phenotypes and failure to find orthologues of *C. elegans sid-2* in previous searches [11,12], our results show that *sid-2* is present in syntenic regions of clade V parasite genomes. The highest conservation is seen in the transmembrane and cytoplasmic domains, and our analysis highlights the divergence of the extracellular domain of SID-2, which shows very little conservation across the clade V nematode orthologues examined. A previous mutagenesis study pinpointed a requirement of the SID-2 extracellular domain for internalisation of dsRNA from the environment: the extracellular domain of *C. elegans* SID-2 was able to rescue an environmental RNAi phenotype in a *sid-2 null* mutant, whereas the divergent extracellular domain from *C. briggsae* SID-2, a species that does not exhibit an environmental RNAi phenotype, does not rescue the phenotype [5]. The high divergence of the extracellular domain may explain the multiple gains and losses of environmental RNAi phenotypes in the *Caenorhabditis* genus, despite all of these species encoding *sid-2* orthologues [6]. The lack of conservation of the extracellular domain is consistent with our finding here that *H. bakeri* SID-2 does not rescue RNAi in *C. elegans* and is unlikely to operate in dsRNA import in the parasite (where environmental RNAi by feeding or soaking in dsRNA has not been shown to work [10]).

The mechanism of dsRNA import by SID-2, and the role of the extracellular domain in this process, is not yet fully understood, but it is proposed to function by pH-dependent receptor-mediated endocytosis [5]. In this regard, it is expected that the extracellular domain of SID-2 acts as a receptor, whereas the cytoplasmic domain interacts with downstream partners in the endocytosis pathway. Here we show that the cytoplasmic domain is relatively conserved across the nematodes which could indicate a common function in endocytosis, but we do not know what molecules / substrates might be internalised by SID-2 In addition to potential variation in the substrate interactions of SID-2 orthologues, a key implication from this work is that the function of SID-2 could extend well beyond the apical membrane of the intestine in the cells where it is synthesised. Our proteomic and western blot analysis shows that SID-2 is highly abundant in extracellular vesicles released from the parasite, which we have previously shown interact with mammalian host cells [15–17]. Recent reports have also demonstrated that some EVs released from *C. elegans* contain SID-2 [13]. An open question is whether the extracellular domain of SID-2 might play a role in the recognition and uptake of EVs by cells and whether this involves RNA. Together these studies expand the contexts in which we should consider SID-2 and its potential interaction partners, and how they may function in the interactions between nematodes and their environments.

## CONCLUSIONS

We identified an orthologue of *C. elegans sid-2* that is highly enriched in *H. bakeri* EVs. Using genome synteny and protein structure-based comparisons, we identified *sid-2* orthologues in other clade V parasitic nematodes. A ConSurf phylogenetic analysis showed that the extracellular domain of SID-2 orthologues in clade V nematodes, which is essential for the environmental RNAi phenotype in *C. elegans*, is more divergent than the transmembrane and cytoplasmic domains. Consistent with previous findings that *H. bakeri* does not exhibit an environmental RNAi phenotype *in vitro*, transgenic expression of *H. bakeri* SID-2 in a *C. elegans* null mutant did not compensate for *C. elegans* SID-2 function in environmental RNAi. We speculate that an ancestral function of nematode SID-2 proteins could relate to EV-mediated communication in free-living and parasitic nematodes.

## METHODS

### *Sid-2* orthologue analysis

Genomic locations and gene accessions of *H. bakeri* and *C. elegans sid-2* orthologues are listed in **Table 1**, and the corresponding amino acid sequences are available in **Table S2**. Signal peptides and transmembrane domains were predicted using DeepTMHMM with default settings [32]. AlphaFold structures were predicted using AlphaFold 3 with default settings [23]. N-glycosylation sites were predicted using NetNGlyC version 1.0 with default settings [33].

Orthologues of *H. bakeri sid-2* HPOL_0001199201 were identified in 14 species of clade V parasitic nematodes using the ‘Orthologues’ function in WormBase ParaSite [34] and manually verified to be nested within a *dyf-2* orthologue. Truncated sequences with length less than 40% of HPOL_0001199201 (358 aa) were excluded from further analyses. We used the protein-to-genome alignment tool miniprot version 0.13 [27] to identify orthologues of HPOL_0001199201 in the genomes of 3 further species of clade V parasitic nematode that are not currently available in WormBase ParaSite: *Teladorsagia circumcincta*, *Trichostrongylus colubriformis*, and *Heligmosomoides polygyrus* (genome accessions are listed in **Table S2**). *Caenorhabditis* nematode orthologues of *C. elegans sid-2* (WBGene00004796) were identified using the ‘Orthologues’ function in WormBase [21]. Details and amino acid sequences of *sid-2* orthologues from clade V parasitic and *Caenorhabditis* nematodes are listed in **Table S2**.

The conservation score amongst clade V nematodes for each amino acid residue in *H. bakeri* SID-2 (HPOL_0001199201) was calculated using the Bayesian method implemented in the ConSurf server [28,29]. A custom multiple sequence alignment (**File S1**) was generated using Clustal Omega with default settings [35] and manually inspected. ConSurf analysis was performed twice to identify regions of HPOL_0001199201 that are conserved with *Caenorhabditis* and other parasitic clade V nematodes: firstly, with all 28 single-copy *sid-2* sequences detailed in **Table S2**; secondly, with only the 13 *sid-2* sequences from parasites detailed in **Table S2**. ConSurf conservation scores were plotted in R Studio [36] using R version 4.4.0 [37] with the packages ggplot2 [38], ggforce [39], and tidyr [40]. The Neighbor-Joining tree generated by ConSurf was visualized in R using the packages ggtree version 3.12.0 [41] and treeio version 1.28.0 [42] (**Fig. S2A**), and the *H. bakeri* Alpha Fold model was visualised with ConSurf conservation scores using PyMol (Schrödinger LLC) (**Fig. S2B**).

### *H. bakeri* extracellular vesicle purification

*H. bakeri* EVs were purified from adult *H. bakeri* excretory-secretory products (HES) collected up to 8 days post-*in vitro* culture as described previously [17]. Eggs and debris were removed from HES by spinning at 400 *x g* for 5 mins at RT and then filtered using a 0.22 μm filter. Filtered HES was concentrated using a VivaSpin 20 centrifugal concentrator with a 5 kDa molecular weight cutoff (Sartorius). EVs were purified from concentrated HES by ultracentrifugation at 100,000 *x g* for 90 mins at 4 °C in polyallomer tubes (Beckman Coulter) in a SW40 rotor (Beckman Coulter). The supernatant (EV-depleted HES) was removed and concentrated using a VivaSpin 20 centrifugal concentrator with a 5 kDa molecular weight cutoff (Sartorius). Purified EVs were washed twice with PBS (Sigma-Aldrich) and pelleted each time by ultracentrifugation at 100,000 *x g* for 70 mins at 4 °C in polyallomer tubes (Beckman Coulter) in a SW40 rotor (Beckman Coulter). The pellet was resuspended in PBS and EV particle size and counts were measured using a Zetaview TWIN particle tracking analyser (Particle Metrix), and protein concentrations were measured using the Qubit Protein Assay (Qubit). EVs were aliquoted and stored at −80 °C until use.

### Silver stain

Samples containing 1 μg total protein were diluted in 1 x loading dye with 0.1 M dithiothreitol and incubated at 70 °C with shaking at 1000 rpm for 10 mins and then separated by electrophoresis on a 4-12% NuPAGE Bis-Tris gel (Invitrogen) at 120 V for 1 h 35 mins. All subsequent steps were performed on a rocker at RT in 100 mL of each solution. Gels were incubated with fixing solution (40% ethanol, 10% glacial acetic acid) for 2 h 30 mins, fixing solution was removed, and gels were incubated with sensitisation solution (30% ethanol, 0.2% sodium thiosulphate, 6.8% sodium acetate) for 30 mins. Gels were washed three times with distilled water for 5 mins and incubated with 0.25% silver nitrate solution for 20 mins. Gels were washed twice with distilled water for 1 min and bands were developed with developing solution (2.5 % sodium carbonate, 20 μl 37% formaldehyde) for 2 mins. Gels were washed once with distilled water for 30 secs and incubated with stopping solution (50 mM EDTA) for 10 mins. Gels were imaged using a ChemiDoc gel imager (BioRad).

### Western blot

Samples containing 1 μg total protein were diluted in 1 x loading dye with 0.1 M dithiothreitol and incubated at 70 °C with shaking at 1000 rpm for 10 mins and then separated by electrophoresis on a 4-12% NuPAGE Bis-Tris gel (Invitrogen) at 120 V for 1 h 40 mins. Proteins were transferred to a PVDF membrane at 100 V for 1 h 45 mins. The membrane was rinsed in Tris-buffered saline with 0.1% Tween-20 (TBS-T) and blocked with 3% milk TBS-T for 1 h 45 mins at RT. Polyclonal antibodies raised in rabbits against the *H. bakeri* SID-2 extracellular domain peptide CSNRVPSGQDDKNITVT (Sino Biological) were diluted 1:1000 in 3% milk TBS-T and incubated with the membrane overnight at 4 °C. Polyclonal antibodies were raised in rats against EVs as detailed in Coakley et al. [16], and for Western blots diluted 1:1000 in 3% milk TBS-T and incubated with the membrane overnight at 4 °C. The membrane was washed four times with TBS-T for 10 mins at RT and then incubated with Goat anti-Rabbit IgG (H+L) Secondary Antibody, DyLight™ 800 4X PEG (Invitrogen, SA5-35571) for anti-SID-2 blots, or Goat anti-Rat IgG (H+L) Cross-Adsorbed Secondary Antibody, Alexa Fluor™ 680 (Life Technologies, A-21096) for anti-EV serum blots, diluted 1:10,000 in 3% milk TBS-T for 1 h at RT. The membrane was washed four times with TBS-T for 10 mins at RT, once with TBS for 5 mins at RT, and then visualised on a LiCor Odyssey imager (LiCor).

### *C. elegans* strains

The *C. elegans* strains were grown on Nematode Growth Media (NGM) plates seeded with *E. coli* OP50 bacteria at 20°C. All strains used in the study are listed in **Table S3**.

### *C. elegans* transgenic strain construction

The Mos1 mediated Single Copy Insertion MosSCI method [43] was used to generate transgenic animals stably expressing (CeSID-2 and HbSID-2) transgenes under the control of the *cesid-2* promoter and *unc-54* 3′ UTR. The transgenes were cloned into the pCFJ151 vector using Gibson assembly [44] and inserted into the chromosome II (*ttTi5605*) locus. Briefly, a mix of plasmids that contains the transgene of interest including a positive selection marker, a transposase plasmid, three fluorescent markers for negative selection [pCFJ90 (Pmyo-2::mCherry), pCFJ104 (Pmyo-3::mCherry) and pGH8 (Prab-3::mCherry)] were injected into *unc- 119* mutant animals. Moving, non-fluorescent worms were then selected, and insertions were confirmed by PCR using primers spanning both homology arms. Both CeSID-2 and HbSID-2 expressing transgenic strains were then crossed into the sid-2 null strain background (PT3646) using standard genetic methods.

### RNAi assay

To perform RNAi-mediated depletion, we designed the targeting sequence for *hcp-4* to be at nucleotide positions 967-2128 after the first ATG codon, as described in Taylor et al. [31]. The *hcp-4* targeting sequence was inserted into the L4440 plasmid and transformed into HT115 (*DE3*) bacteria [45]. Bacterial clones containing the RNAi sequence were cultured overnight at 37°C in LB medium with 100 μg mL^-1^ ampicillin. Saturated cultures were diluted 1:100 and grown until reaching an OD600 of 0.8-1. Isopropyl-β-D-thiogalactopyranoside (IPTG) was added to a final concentration of 1 mM, and the cultures were incubated for 1 hour at 37°C. The bacteria were then seeded onto NGM plates containing agarose and 1 mM IPTG, and the plates were allowed to dry. L4 worms were subsequently plated on RNAi plates, maintained at 20°C and the RNAi assay was performed as outlined in Fig. 4A and as follows:

On experimental Day 1, a single L4 hermaphrodite of the desired strain was placed onto each seeded plate (n=10). After 24 hours (Day 2), the same animal (now an adult) was transferred to a fresh seeded plate. This process was repeated every 24 hours, moving the animal to a new plate on Day 3 and removing it on Day 4. Following the removal of the animal, each plate was incubated at 20°C for 48 hours before counting the number of live L4 progeny and dead/unhatched eggs. The number of dead eggs was added to the number of live offspring for each plate, and the percentage viability of the strain after *hcp-4* dsRNA ingestion was determined by calculating the number of live progeny divided by the total progeny. Statistical analyses were performed in GraphPad prism version 10.2.3 (GraphPad Software) with a significance α threshold of 0.05. Due to heterogeneity of variance between groups, percentage viability of offspring in the RNAi assay was analyzed using a Kruskal-Wallis test followed by Dunn’s test for post-hoc comparisons between groups.

### *C. elegans* live imaging

For all imaging experiments, L4 animals were anesthetised using 5 mM levamisole and mounted in M9 on 2% agarose pads. Images were acquired with a CFI60 Plan Apochromat lambda 100X (Nikon) objective mounted on a spinning disc confocal microscopy system. The system was equipped with a Yokogawa spinning disk unit (CSU-W1), a Nikon Ti2-E fully motorised inverted microscope, and a Photometrics Prime 95B camera.

To image the localisation of CeSID-2::GFP and HbSID-2::GFP in the gut, a single z-slice of the gut was focused at the centre for still images. For time-lapse movies, a single z-slice of the apical surface of the gut was captured every two seconds for a total duration of 2 minutes. All acquired images were then processed using ImageJ (Fiji) software.

## SUPPLEMENTARY

**Table S1** Twenty most abundant proteins identified by LC-MS/MS of *H. bakeri* extracellular vesicle fractions of worm secretion products (Buck et al. 2014 Dataset S2 re-annotated with gene identifiers for genome PRJEB15396 from WormBase ParaSite) and their annotated functions. HPOL_0001199201 (highlighted in grey) is the nineteenth most abundant protein.

**Table S2** Amino acid sequences identified as *sid-2* orthologues in *Caenorhabditis* and parasitic clade V nematodes. All *sid-2* orthologues were single-copy in the genomes examined. Splice variants and sequences that were shorter than 40% length of HPOL_0001199201 (highlighted in grey) were excluded from ConSurf analysis.

**Table S3** *C. elegans* strains used in this study.

**Figure S1** Protein sequence and structural features of the *H. bakeri sid-2* orthologue that is enriched in EVs compared to EV-depleted HES. **A** Amino acid alignment showing low sequence conservation between *H. bakeri* and *C. elegans sid-2* orthologues (24.5 % sequence identity). Regions predicted by DeepTMHMM [32] to be signal peptide, cytoplasmic domain, transmembrane helix and extracellular domain are annotated for each sequence and detailed in the table on the right. AlphaFold 3 predicted structures for *sid-2* orthologues from **B** *H. bakeri* and **C** *C. elegans*. Structures are coloured by the pLDDT confidence estimate, which is scored from 0-100 (100 = higher confidence) and color coded as follows: dark blue is very high (pLDDT > 90), cyan is confident (90 > pLDDT > 70), yellow is low (70 > pLDDT > 50), and orange is very low (pLDDT < 50). Predicted aligned error is shown for each structure, with dark green indicating likely proximity of residues.

**Figure S2** ConSurf analysis of Clade V nematode *sid-2* orthologues amino acid sequences **A** Neighbor-Joining phylogeny generated by ConSurf analysis, *C. elegans* and *H. bakeri* sequences are highlighted in grey. **B** *H. bakeri* SID-2 AlphaFold model with residues coloured by ConSurf conservation scores from analysis of all sequences (top) and parasite sequences only (bottom).

**File S1** Multiple sequence alignment of all amino acid sequences used for ConSurf analysis.

**Movie S1** Movement of transgenic GFP in the intestinal cells of transgenic *C. elegans* strain DKC1365 expressing *C. elegans* SID-2 indicates the association of SID-2 transgenes with vesicles in the intestine.

**Movie S2** Movement of transgenic GFP in the intestinal cells of transgenic *C. elegans* strain DKC1285 expressing *H. bakeri* SID-2 indicates the association of SID-2 transgenes with vesicles in the intestine.

## ACKNOWLEDGEMENTS

We thank Elaine Robertson for generating parasite excretory-secretory material, Kyriaki Neophytou and Lewis Strachan for initial testing of the SID-2 antibody, Lilli Skäer for preliminary experiments, and Ruby White and Elaine Robertson for generating anti-EV serum. We thank Cameron Finlayson for help with *C. elegans* rearing, and Cei Abreu-Goodger and Isaac Martínez-Ugalde for advice on orthologue finding. FB and AHB are supported by ERC Consolidator Award 101002385, D.K.C. is supported by a Sir Henry Dale Fellowship jointly funded by the Wellcome Trust and the Royal Society (208833/Z/17/Z) and BP by a Sir Henry Wellcome Postdoctoral Fellowship (215925). Imaging was done at the microscopy facility at the the Wellcome Centre for Cell Biology funded by the core grant 203149 and Wellcome Discovery Research Platform Award 226791.

## ETHICS

Research carried out at the University of Edinburgh is subject to review by the School of Biological Sciences ethics committee and maintenance of the parasite life cycle and generation of parasite excretory-secretory products is conducted in accordance with the UK Home Office and local ethically approved guidelines.

## USE OF ARTIFICIAL INTELLIGENCE AND AI-ASSISTED TECHNOLOGIES

We used the software AlphaFold 3, which employs AI to predict protein 3D structure.

## DATA, CODE AND MATERIALS

The datasets supporting this article have been uploaded as part of the supplementary material.

## AUTHORS’ CONTRIBUTIONS

FB: Conceptualization, data curation, formal analysis, investigation, methodology, project administration, software, supervision, validation, visualization, writing-original draft, writing-review and editing

KJ: Data curation, formal analysis, investigation, project administration, visualization, writing-original draft, writing-review and editing

FWNC: Formal analysis, visualization, writing-review and editing

IN: Investigation, methodology, resources, writing-review and editing MBB: Resources, writing-review and editing

AGC: Methodology, writing-review and editing

BP: Methodology, investigation, writing-review and editing

DKC: Conceptualization, data curation, funding acquisition, investigation, methodology, project administration, supervision, visualization, writing-original draft, writing-review and editing

AHB: Conceptualization, funding acquisition, methodology, supervision, writing-original draft, writing-review and editing

## REFERENCES

1. Whangbo JS, Hunter CP. 2008 Environmental RNA interference. Trends Genet. 24, 297–305.

2. Tijsterman M, May RC, Simmer F, Okihara KL, Plasterk RHA. 2004 Genes required for systemic RNA interference in Caenorhabditis elegans. Curr. Biol. 14, 111–116.

3. Winston WM, Molodowitch C, Hunter CP. 2002 Systemic RNAi in C. elegans requires the putative transmembrane protein SID-1. Science 295, 2456–2459.

4. Winston WM, Sutherlin M, Wright AJ, Feinberg EH, Hunter CP. 2007 Caenorhabditis elegans SID-2 is required for environmental RNA interference. Proc. Natl. Acad. Sci. U. S. A. 104, 10565–10570.

5. McEwan DL, Weisman AS, Hunter CP. 2012 Uptake of extracellular double-stranded RNA by SID-2. Mol. Cell 47, 746–754.

6. Nuez I, Félix M-A. 2012 Evolution of susceptibility to ingested double-stranded RNAs in Caenorhabditis nematodes. PLoS One 7, e29811.

7. Blaxter ML et al. 1998 A molecular evolutionary framework for the phylum Nematoda. Nature 392, 71–75.

8. Smythe AB, Holovachov O, Kocot KM. 2019 Improved phylogenomic sampling of free-living nematodes enhances resolution of higher-level nematode phylogeny. BMC Evol. Biol. 19, 121.

9. Maizels RM et al. 2012 Immune modulation and modulators in Heligmosomoides polygyrus infection. Exp. Parasitol. 132, 76–89.

10. Lendner M, Doligalska M, Lucius R, Hartmann S. 2008 Attempts to establish RNA interference in the parasitic nematode Heligmosomoides polygyrus. Mol. Biochem. Parasitol. 161, 21–31.

11. Viney ME, Thompson FJ. 2008 Two hypotheses to explain why RNA interference does not work in animal parasitic nematodes. Int. J. Parasitol. 38, 43–47.

12. Dalzell JJ et al. 2011 RNAi effector diversity in nematodes. PLoS Negl. Trop. Dis. 5, e1176.

13. Nikonorova IA et al. 2022 Isolation, profiling, and tracking of extracellular vesicle cargo in Caenorhabditis elegans. Curr. Biol. 32, 1924–1936.e6.

14. Wang J, Silva M, Haas LA, Morsci NS, Nguyen KCQ, Hall DH, Barr MM. 2014 C. elegans ciliated sensory neurons release extracellular vesicles that function in animal communication. Curr. Biol. 24, 519–525.

15. Buck AH et al. 2014 Exosomes secreted by nematode parasites transfer small RNAs to mammalian cells and modulate innate immunity. Nat. Commun. 5, 5488.

16. Coakley G et al. 2017 Extracellular Vesicles from a Helminth Parasite Suppress Macrophage Activation and Constitute an Effective Vaccine for Protective Immunity. Cell Rep. 19, 1545–1557.

17. Chow FW-N et al. 2019 Secretion of an Argonaute protein by a parasitic nematode and the evolution of its siRNA guides. Nucleic Acids Res. 47, 3594–3606.

18. White R et al. 2023 Special considerations for studies of extracellular vesicles from parasitic helminths: A community-led roadmap to increase rigour and reproducibility. J Extracell Vesicles 12, e12298.

19. Altschul SF, Gish W, Miller W, Myers EW, Lipman DJ. 1990 Basic local alignment search tool. J. Mol. Biol. 215, 403–410.

20. Stevens L et al. 2023 Ancient diversity in host-parasite interaction genes in a model parasitic nematode. Nat. Commun. 14, 7776.

21. Davis P et al. 2022 WormBase in 2022-data, processes, and tools for analyzing Caenorhabditis elegans. Genetics 220. (doi:10.1093/genetics/iyac003)

22. Carlton PM, Davis RE, Ahmed S. 2022 Nematode chromosomes. Genetics 221. (doi:10.1093/genetics/iyac014)

23. Abramson J et al. 2024 Accurate structure prediction of biomolecular interactions with AlphaFold 3. Nature (doi:10.1038/s41586-024-07487-w)

24. van Kempen M, Kim SS, Tumescheit C, Mirdita M, Lee J, Gilchrist CLM, Söding J, Steinegger M. 2024 Fast and accurate protein structure search with Foldseek. Nat. Biotechnol. 42, 243–246.

25. Rath A, Glibowicka M, Nadeau VG, Chen G, Deber CM. 2009 Detergent binding explains anomalous SDS-PAGE migration of membrane proteins. Proc. Natl. Acad. Sci. U. S. A. 106, 1760–1765.

26. Magnelli PE, Bielik AM, Guthrie EP. 2011 Identification and characterization of protein glycosylation using specific endo- and exoglycosidases. *J. Vis. Exp.* , e3749.

27. Li H. 2023 Protein-to-genome alignment with miniprot. Bioinformatics 39. (doi:10.1093/bioinformatics/btad014)

28. Ashkenazy H, Erez E, Martz E, Pupko T, Ben-Tal N. 2010 ConSurf 2010: calculating evolutionary conservation in sequence and structure of proteins and nucleic acids. Nucleic Acids Res. 38, W529–33.

29. Ashkenazy H, Abadi S, Martz E, Chay O, Mayrose I, Pupko T, Ben-Tal N. 2016 ConSurf 2016: an improved methodology to estimate and visualize evolutionary conservation in macromolecules. Nucleic Acids Res. 44, W344–50.

30. Moore LL, Roth MB. 2001 HCP-4, a CENP-C-like protein in Caenorhabditis elegans, is required for resolution of sister centromeres. J. Cell Biol. 153, 1199–1208.

31. Taylor SJP, Bel Borja L, Soubigou F, Houston J, Cheerambathur DK, Pelisch F. 2023 BUB-1 and CENP-C recruit PLK-1 to control chromosome alignment and segregation during meiosis I in C. elegans oocytes. Elife 12. (doi:10.7554/eLife.84057)

32. Hallgren J, Tsirigos KD, Pedersen MD, Armenteros JJA, Marcatili P, Nielsen H, Krogh A, Winther O. 2022 DeepTMHMM predicts alpha and beta transmembrane proteins using deep neural networks. bioRxiv. , 2022.04.08.487609. (doi:10.1101/2022.04.08.487609)

33. Gupta R, Brunak S. 2002 Prediction of glycosylation across the human proteome and the correlation to protein function. Pac. Symp. Biocomput. , 310–322.

34. Howe KL, Bolt BJ, Shafie M, Kersey P, Berriman M. 2017 WormBase ParaSite -a comprehensive resource for helminth genomics. Mol. Biochem. Parasitol. 215, 2–10.

35. Madeira F, Pearce M, Tivey ARN, Basutkar P, Lee J, Edbali O, Madhusoodanan N, Kolesnikov A, Lopez R. 2022 Search and sequence analysis tools services from EMBL-EBI in 2022. Nucleic Acids Res. 50, W276–W279.

36. RStudio Team. 2021 RStudio: Integrated Development Environment for R.

37. R Core Team. 2024 R: A Language and Environment for Statistical Computing.

38. Wickham H. 2016 ggplot2: Elegant Graphics for Data Analysis.

39. Pedersen TL. 2024 ggforce: Accelerating “ggplot2.”

40. Wickham H, Vaughan D, Girlich M. 2024 tidyr: Tidy Messy Data.

41. Yu G, Smith DK, Zhu H, Guan Y, Lam TT-Y. 2017 Ggtree: An r package for visualization and annotation of phylogenetic trees with their covariates and other associated data. Methods Ecol. Evol. 8, 28–36.

42. Wang L-G et al. 2020 Treeio: An R Package for Phylogenetic Tree Input and Output with Richly Annotated and Associated Data. Mol. Biol. Evol. 37, 599–603.

43. Frøkjaer-Jensen C, Davis MW, Hopkins CE, Newman BJ, Thummel JM, Olesen S-P, Grunnet M, Jorgensen EM. 2008 Single-copy insertion of transgenes in Caenorhabditis elegans. Nat. Genet. 40, 1375–1383.

44. Gibson DG, Young L, Chuang R-Y, Venter JC, Hutchison CA 3rd, Smith HO. 2009 Enzymatic assembly of DNA molecules up to several hundred kilobases. Nat. Methods 6, 343–345.

45. Timmons L, Court DL, Fire A. 2001 Ingestion of bacterially expressed dsRNAs can produce specific and potent genetic interference in Caenorhabditis elegans. Gene 263, 103–112.

