## Supplementary material for "SID-2 localises to extracellular vesicles in parasitic nematodes and does not function in environmental RNAi": Fig S1 S2 Table S1 S3

## A

```
                       Signal peptide              Extracellular
H. bakeri     1 MPS-WRHL-LLLVVIVDVTSAGLKNVVVVQNFNL------KTAVAAEISCSNIKSVLVEG
C. elegans    1 MPR-FVYFCFALIALL-PISWT-MDGILITDVEI------HV-DVCQISCKASNTASLLI

H. bakeri    53 RAV-GSYSCAGATTTTNNTQSGMWIVVETQT------------------------V-IPK
C. elegans   51 NDAPFTPMCNSAGD--------QIFFTYNGT------------------------AAISD

H. bakeri    87 DKTVSWRIEYTTGY----------------LASSFTVDKALYKPLSSTVSLPVSISTKAL
C. elegans   79 LKNVTFILEVTTDTK--------NCTFTANYTGYFTPDPKSK----------------PF

H. bakeri   131 SLLSNRVPSGQDDKNITVTTIASSEITTLKPTTESKAPEPSKAPTKP--PTSEAPTP---
C. elegans  115 QLGFASATLNRD-----------MGKVTKTI--------MEDSG-----EMVEQDFSNSS
                                                          Transmembrane helix
H. bakeri   186 ----EP--TTKPTTVAVNVTVGYIQFQTGKE--NL-QNQYTQAVVTAVIEGLLLGAILFV
C. elegans  151 AVPTPASTTPLPQSTVAHLTIAYVHLQYEE-TKTV-VNKNGGAVAVAVIEGIALIAILAF
                    Cytoplasmic
H. bakeri   237 FLFKCYQRSKLRTAGMYPTRNDFQLNASSRNTVFYDNGGTLSYG--AGPNARPEIPSYRS
C. elegans  209 LGYRTMVNHKLQNSTRTNGLYGYDNNNSSRITVPDAMR--M------SDIPPPRDPMYAS

H. bakeri   295 QDSQ------PPLRLDSLTPRPAQQV-VTPTLMPDSSRTLSPPVT-TVQP----------
C. elegans  261 PPTP-----------LSQPTPARNTVMTTQELVVPTA-NSSAAQP-ST-------TSNGQ

H. bakeri   337 -----APLNYWPDQP---QPQPVKNIMDSDL
C. elegans  301 FNDPFATLESW--------------------
```

| | *H. bakeri* | | *C. elegans* | |
|---|---|---|---|---|
| | **Start aa** | **End aa** | **Start aa** | **End aa** |
| Signal peptide | 1 | 19 | 1 | 20 |
| Extracellular domain | 20 | 218 | 21 | 193 |
| Transmembrane helix | 219 | 239 | 194 | 211 |
| Cytoplasmic domain | 240 | 358 | 212 | 311 |

## B

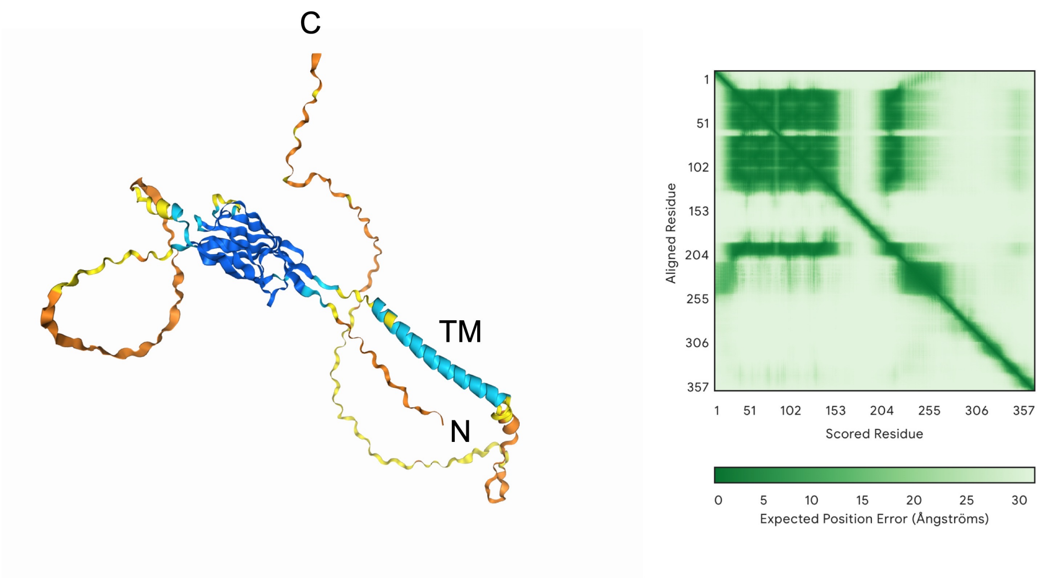


## C

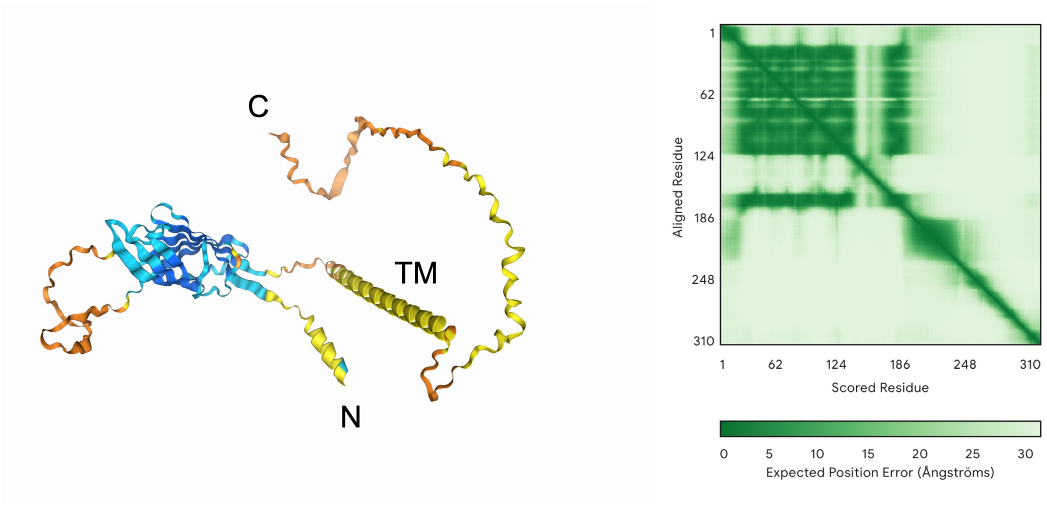

### Fig. S2

**A**

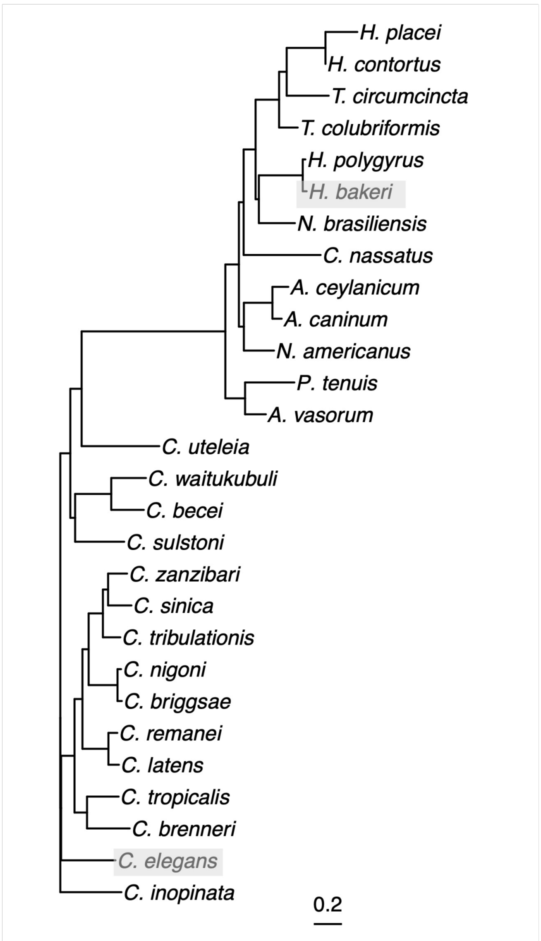
H. placei
H. contortus
T. circumcincta
T. colubriformis
H. polygyrus
H. bakeri
N. brasiliensis
C. nassatus
A. ceylanicum
A. caninum
N. americanus
P. tenuis
A. vasorum
C. uteleia
C. waitukubuli
C. becei
C. sulstoni
C. zanzibari
C. sinica
C. tribulationis
C. nigoni
C. briggsae
C. remanei
C. latens
C. tropicalis
C. brenneri
C. elegans
C. inopinata
0.2

**B**

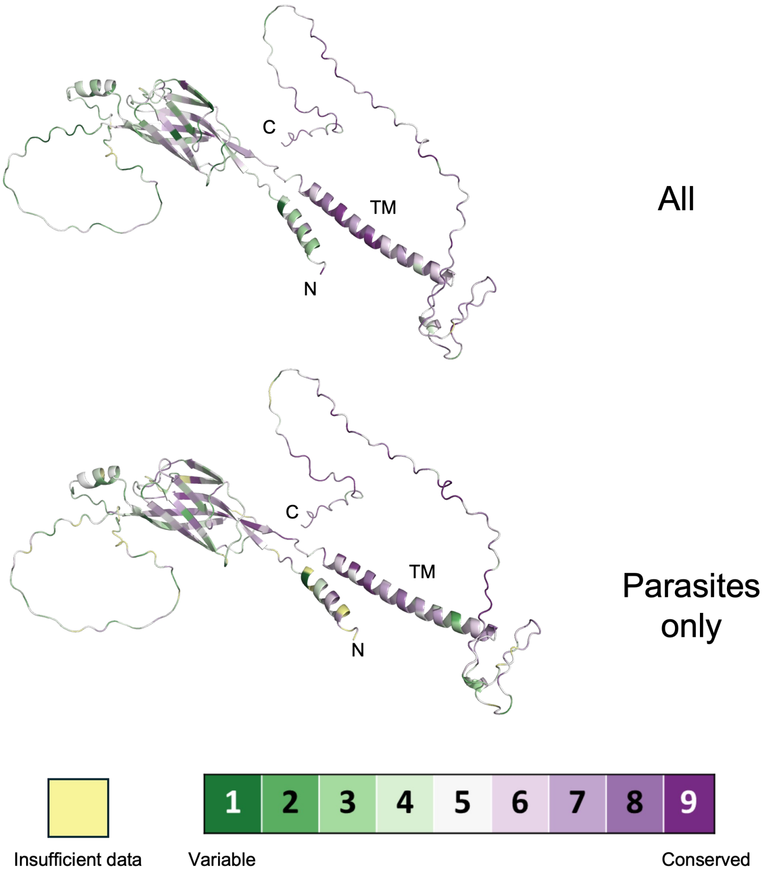
C
TM
N
All
C
TM
N
Parasites only
1
2
3
4
5
6
7
8
9
Insufficient data
Variable
Conserved

### Table S1

| Rank | Accession | Pfam annotated functional domain description | Pfam accession |
|---|---|---|---|
| 1 | HPOL_0001315501 | Peptidase M13, C-terminal domain; Peptidase M13, N-terminal domain | PF01431; PF05649 |
| 2 | HPOL_0001276401 | Peptidase M13, C-terminal domain; Peptidase M13, N-terminal domain | PF01431; PF05649 |
| 3 | HPOL_0001047201 | Chitin binding domain superfamily | PF01607 |
| 4 | HPOL_0001695101 | Peptidase M13, C-terminal domain; Peptidase M13, N-terminal domain | PF01431; PF05649 |
| 5 | HPOL_0000148201 | Ezrin/radixin/moesin, C-terminal; Ezrin/radixin/moesin, alpha-helical domain; FERM central domain; FERM, C-terminal PH-like domain; FERM, N-terminal | PF00769; PF20492; PF00373; PF09380; PF09379 |
| 6 | HPOL_0002004401 | Peptidase family A1 domain | PF00026 |
| 7 | HPOL_0001109801 | - | |
| 8 | HPOL_0002023901 | Vitellinogen, open beta-sheet; Vitellogenin, N-terminal; von Willebrand factor, type D domain | PF09172; PF01347; PF00094 |
| 9 | HPOL_0001496301 | Aminopeptidase N-like, N-terminal domain; ERAP1-like C-terminal domain; Peptidase M1, membrane alanine aminopeptidase | PF17900; PF11838; PF01433 |
| 10 | HPOL_0001549601 | SnoaL-like domain | PF13474 |
| 11 | HPOL_0001502601 | Peptidase family A1 domain | PF00026 |
| 12 | HPOL_0001631701 | Annexin repeat | PF00191 |
| 13 | HPOL_0002135201 | - | - |
| 14 | HPOL_0001876901 | Heat shock protein 70 family | PF00012 |
| 15 | HPOL_0000529301 | - | - |
| 16 | HPOL_0001611801 | - | - |
| 17 | HPOL_0001832701 | - | - |
| 18 | HPOL_0000312901 | Ubiquitin-like domain | PF00240 |
| 19 | HPOL_0001199201 | - | - |
| 20 | HPOL_0001165701 | Vitellogenin, N-terminal | PF01347 |

### Table S3

| Genotype | Source | Identifier |
|---|---|---|
| *C. elegans N2 Bristol* | Caenorhabditis Genetics Center | N2 |
| *sid-2(my99[Q163stop]) III; myIs4 [pkd-2::GFP + ccGFP]; him-5 (e1490) V* | Nikonorova et al. 2022 | PT3646 |
| *dhaSi320[pDC1210; Cesid-2pro::Hbsid-2::GFP::unc-54 3'UTR; cb-unc-119(+)] II (clone B3); sid-2(my99[Q163stop]) III* | This study | DKC1285 |
| *dhaSi329[pDC1223; Cesid-2pro::Cesid-2::GFP::unc-54 3'UTR; cb-unc-119(+)]; sid-2(my99[Q163stop]) III* | This study | DKC1365 |
